# Cytidine deaminase regulates mitochondrial biogenesis in pancreatic cancer cells

**DOI:** 10.1101/2022.12.15.520429

**Authors:** Audrey Frances, Audrey Lumeau, Lucille Stuani, Marion Gayral, Estelle Saland, Nicolas Bery, Delphine Pagan, Naima Hanoun, Jérôme Torrisani, Anthony Lemarié, Jean-Charles Portais, Louis Buscail, Nelson Dusetti, Jean-Emmanuel Sarry, Pierre Cordelier

## Abstract

Despite tremendous efforts from the scientific community, pancreatic ductal adenocarcinoma (PDAC) is still a deadly disease and will soon become the second cause of death by cancer worldwide^1^. When surgery is not possible, therapeutic options are few and ineffective^2^. Patients are most often treated with Folfirinox or gemcitabine chemotherapy, that extends survival in weeks to months. Cytidine deaminase (CDA) catalyzes the irreversible hydrolytic deamination of cytidine and deoxycytidine to uridine and deoxyuridine to fuel RNA and DNA synthesis^3^. CDA also deaminates and neutralizes deoxycytidine-based therapies, and as such, has been identified as a major contributor of tumor chemoresistance, especially to gemcitabine in PDAC^3^. We previously identified that CDA is elevated in PDAC tumors at diagnosis and that CDA exerts an unexpected role on DNA replication that can be exploited for therapeutic intervention^4^. Very recently, CDA was associated with cellular metabolism^5^. Here, we show that CDA promotes mitochondrial biogenesis and oxidative phosphorylation independently of its deaminase activity. This uncloaks novel therapeutic vulnerabilities in primary cancer cells that overexpress this protein. This study shines a new light on the tumoral potential of CDA in PDAC.

## RESULTS

To fully capture the role of CDA on PDAC cells metabolism, we overexpressed wild-type (wt) and catalytically-inactive CDA (E67Q) using lentiviral vectors in human Mia PaCa-2 cells that fully recapitulate genetic alterations found in PDAC tumors, and that express moderate levels of CDA as compared to other PDAC cell lines^4^. Control cells expressed luciferase. Alternatively, we generated Mia Paca-2 cells expressing control shRNA, or shRNA targeting CDA^4^. Cell lines were validated^4^, and we performed transcriptomic studies to identify that CDAE67Q expression resulted in the enrichment of signatures related to mitochondria biology and activity (oxidative phosphorylation, normalized enrichment score (NES)=2.2, adjusted *p* value=2.8e-9, mitochondrial translation, NES=2.2, adjusted *p* value=1.7e-7, mitochondrial gene expression, NES=1.9, adjusted *p* value=1.4e5, electron transport chain, NES=1.6, adjusted *p* value=1.6e6). We found similar transcriptomic enrichment signature in PDAC tumors from the TCGA with high expression of CDA (mitochondrial translation, NES=1.9, adjusted *p* value=8e5, oxidative phosphorylation, NES=2.1, adjusted *p* value=1.3e6). We further analysed the mitochondrial landscape of cells and found that CDA and CDAE67Q significantly increases mitochondrial mass (+18%±8% and +39%±11%, respectively, *p*<0.05, Figure 1A). On the contrary, we found a significant reduction in mitochondrial mass (−20%±4%, *p*<0.05, Figure 1B), in mitochondria number (− 21%±11%, *p*<0.05, Figure 1C and Supplemental Figure 1A), surface (−33%±10%, *p*<0.0001, Supplemental Figure 1B) and width (−23%±2%, *p*<0.0001, Supplemental Figure 1C) in cells expressing CDA shRNA as compared to control cells. CDA and CDAE67Q expression significantly increased mitochondrial DNA content in PDAC cells (+35%±8% and +24%±5%, respectively, *p*<0.05, Figure 1D), and the expression of mitochondrial transcription factor A (TFAM, a key activator of mitochondrial transcription and mitochondrial genome replication), mitochondrial complex proteins, isocitrate dehydrogenase (NAD(+)) 3 catalytic subunit alpha (IDH3A) and succinate dehydrogenase complex, subunit A (SDHA), as compared to control cells (Figure 1E). By Western blot in PDAC cells expressing CDA at endogenous levels, we found that CDA is not a mitochondrial protein (Supplemental Figure 1D). We further analysed markers of mitochondrial fusion (mitofusin Mfn1, and optic atrophy-1, OPA1, protein) or fission (Dynamin-related protein 1, DRP1) in PDAC cells but found no significant nor consistent changes in response to CDA or CDAE67Q overexpression (Supplemental Figure 1E). Conversely, targeting CDA with shRNA resulted in the reduction of mitochondrial DNA content in PDAC cells (Figure 1D, −35%±14%, *p*<0.05), and significant decrease in the expression of TFAM, mitochondrial complex proteins, IDH3A and SDHA (Figure 1E). These results point towards a role of CDA on mitogenesis rather than on mitochondrial dynamics. Importantly, diazepinone riboside (DR) and tetrahydrouridine (THU), two catalytic inhibitors of CDA, showed no effect on the mitochondrial DNA content and on the expression of IDH3A and SDHA proteins (Supplemental Figure 1F & 1G), contrary to CDA overexpression or downregulation (Figure 1D & 1E).

**Figure 1.**
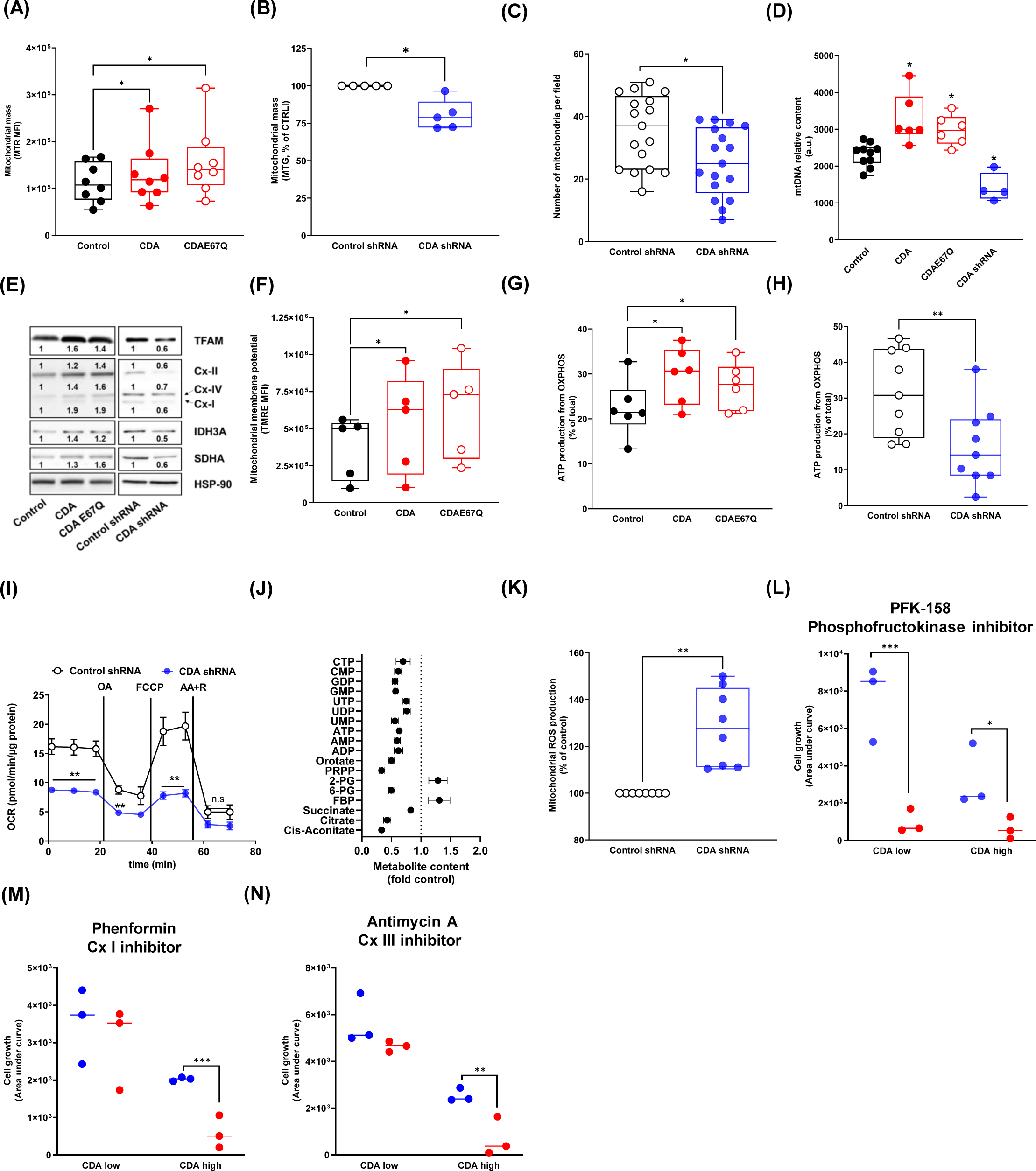
Quantification of mitochondrial mass using mitotracker red (MTR) in Mia PaCa-2 cells overexpressing wtCDA or CDAE67Q. (**A**), or expressing shRNA targeting CDA (**B**), respectively. Cells expressing luciferase or random shRNA were used as control. Results are expressed as mean ± min to max of five to eight independent experiments using two to three different cell preparation analyzed in duplicate. *: *p*<0.05. **C**. Mia PaCa-2 cells expressing shRNA against CDA were analyzed by electron microscopy for mitochondria number enumeration. Cells expressing random shRNA were used as control. Results are expressed as mean ± min to max of two independent experiments using seventeen different fields. *: *p*<0.05. **D**. Mitochondrial DNA (mtDNA) content was analyzed by qPCR, in Mia PaCa-2 cells overexpressing wtCDA, CDAE67Q, or expressing shRNA against CDA. Cells expressing luciferase or random shRNA were used as control. Results are expressed as mean ± min to max of four to eight independent experiments using two or three different cell preparation analyzed in triplicate, respectively. *: *p*<0.05. **E**. Western blotting for TFAM, mitochondrial complex I, III and IV proteins, IDH3A and SDHA in Mia PaCa-2 cells overexpressing wtCDA, CDAE67Q, or expressing shRNA against CDA. Cells expressing luciferase or random shRNA were used as control. HSP90 was used as a loading control. Fold-change in band intensity is indicated. Representative of three independent experiments. **F**. Mitochondrial membrane potential analysis using tetramethylrhodamine, ethyl ester (TMRE) in Mia PaCa-2 cells overexpressing wtCDA or CDAE67Q. Cells expressing luciferase were used as control. Results are expressed as mean ± min to max of five independent experiments using two different cell preparation analyzed in triplicate. *: *p*<0.05. Quantification of ATP production from OXPHOS in Mia PaCa-2 cells overexpressing wtCDA, CDAE67Q (**G**), or expressing shRNA against CDA (**H**). Cells expressing luciferase or random shRNA were used as control. Results are expressed as mean ± min to max of six to nine independent experiments using two or three different cell preparation analyzed in triplicate, respectively. *: *p*<0.05, **: *p*<0.01. I. Oxygen consumption rate measurement in Mia PaCa-2 cells expressing shRNA against CDA. Cells expressing random shRNA were used as control. Results are mean ± s.e.m. of four independent experiments performed in triplicate. **: *p*<0.01. OA:, FCCP: Carbonyl cyanide 4-(trifluoromethoxy)phenylhydrazone, R: rotenone, OA: oligomycine A, AA: antimycin A. **J**. Untargeted metabolomic analysis of Mia PaCa-2 cells expression random shRNA or shRNA targeting CDA. Results are mean± s.e.m of three independent experiments. P value ranging from <0.05 to <0.0001. CTP: cytosine triphosphate, CMP: cytosine monophosphate, GDP: guanosine diphosphate, GMP: guanosine monophosphate, UTP: uridine triphosphate, UDP: uridine diphosphate, UMP: uridine monophosphate, ATP: adenosine triphosphate, AMP: adenosine monophosphate, ADP: adenosine diphosphate, PRPP: phosphor ribosyl pyrophosphate, 2-PG: 2-phosphoglyceric acid, 6-PG: 6-phosphogluconate, FBP: fructose 1,6-bisphosphate. **K**. Mitochondrial reactive oxygen species (ROS) production in Mia PaCa-2 cells expressing shRNA against CDA. Cells expressing random shRNA were used as control. Results are mean ± s.e.m. of eight independent experiments using two different cell preparation. **: *p*<0.01. Primary PDAC cells expressing low or high level of CDA (n=3 each) were cultured in the presence of 50 μm PFK-158 (**L**), 0.5 mM phenformin (**M**) and 10 μM Antimycin A (**N**) for 72 hours. Cell growth was monitored longitudinally and expressed as area under curve (AUC). Results are mean ± s.e.m. of two independent experiment with six repetitions per cell line. *: *p*<0.05, **: *p*<0.01, ***: *p*<0.005. Blue dots: mock-treated cells, red dots: cells treated with the corresponding drug.

Furthermore, functional studies identified that CDA and CDAE67Q expression increases mitochondrial membrane potential in PDAC cells (+42%±22% and +67%±24% respectively, *p*<0.05, Figure 1F). Increased mitochondria activity in cells expressing CDA or CDE67Q results in increased production of ATP from oxidative phosphorylation (OXPHOS, +33%±9% and +22%±9% respectively, *p*<0.05, Figure 1G), while ATP production from OXPHOS was reduced when cells expressed shRNA against CDA (−48%±17%, *p*<0.01, Figure 1H). Here again, catalytic inhibitors of CDA showed no activity on ATP production in PDAC cells (Supplemental Figure 1H), further indicating that CDA exerts its novel effect on mitochondria independently of its known catalytic activity. In PDAC cells expressing shRNA against CDA, basal respiration and maximal respiratory capacity are severely compromised (−47%±20%, *p*<0.01 and −62%±4%, *p*<0.005 respectively, Figure 1I). We performed untargeted metabolomics studies and found that the production of several metabolites from the tricarboxylic acid cycle (citrate, isocitrate, succinate) were impaired in cells expressing shRNA against CDA as compared to control cells (Figure 1J). On the contrary, fructose-1,6-bisphoshate and 2-phosphoglycerate levels were elevated in the cells, suggesting compensatory glycolysis consecutive to CDA targeting (Figure 1J). In addition, the levels of phosphoribosyl pyrophosphate and orotate were significantly diminished in cells depleted for CDA (Figure 1J). This may account for the global decrease in purine and pyrimidine synthesis in these cells (Figure 1J). Last, the production of mitochondrial reactive oxygen species (ROS), that are well-characterized markers of mitochondrial respiratory chain dysfunction, is significantly induced in cells depleted for CDA before cells undergo cell death by apoptosis (+28%±5%, *p*<0.01, Figure 1K).

Collectively, we uncovered that CDA plays an unexpected role in cancer cells as it promotes mitochondrial biogenesis that correlates with increased OXPHOS capacity. We then asked whether this would create a metabolic vulnerability in primary cultures. Thus, we selected two distinct groups of three primary cells derived from patients with PDAC with known CDA expression, that we named CDA high and CDA low (Supplemental Figure 1I and 1J). We challenged these cells with PFK-158, a phosphofructokinase inhibitor, and electron transport chain (ETC) I (phenformin), and III (antimycin A) inhibitors. We found that primary PDAC cells proliferation was strongly inhibited following PFK-158, regardless of CDA expression (−87%±18%, *p*<0.01 for CDA low-cells, and −81%±24% *p*<0.05 for CDA high-cells, respectively, Figure 1L). On the contrary, targeting mitochondrial ETC strongly inhibited PDAC primary cells proliferation with high CDA expression only (−71%±18%, *p*<0.005 for phenformin, and −72%±15% *p*<0.01 for antimycin A, respectively, Figure 1M and 1N).

## DISCUSSION

The importance of mitochondria for the initiation and progression of pancreatic tumorigenesis is now well accepted^6^. However, the molecular mechanisms involved remain largely unknown. Here, we found that CDA, that belongs to the pyrimidine salvage pathway, has a new and unexpected role on mitochondria biology and OXPHOS in PDAC cells. Within the *de novo* pyrimidine synthesis, dihydroorotate dehydrogenase (DHODH) is inserted into the outer face of the mitochondrial inner membrane^7^. DHODH depends on respiration to catalyze the fourth and last step in pyrimidine metabolism, and in turn, participates to OXPHOS with its electron acceptor Coenzyme Q^7^. However, we found that CDA is not a mitochondrial protein (Supplemental Figure 1D). In addition, our results suggest that CDA function on mitochondria and OXPHOS may likely occur independently of the known catalytic activity of the enzyme. Thus, while this study comforts the previously known interconnection between mitochondrial function and pyrimidine production in cells, our work calls for additional molecular investigations to fully capture the role of CDA on mitochondria biogenesis and function in PDAC cells. In addition, we show here that the increase of OXPHOS activity in cells expressing high levels of CDA uncloaks novel therapeutic vulnerabilities for mitochondria targeting-drugs, such as phenformin. Unfortunately, phenformin, that shows antitumor activity in patient-derived xenografts^8^, has been so far disappointing in clinical trials, including in PDAC, due to a general lack of efficacy and safety issues^9^. As phenformin side effects can now be clinically managed^10^, our work provides rationale for considering CDA expression for the selection of patients with PDAC that may benefit the most from this molecule.

## FIGURE LEGENDS

**Supplemental Figure 1.**
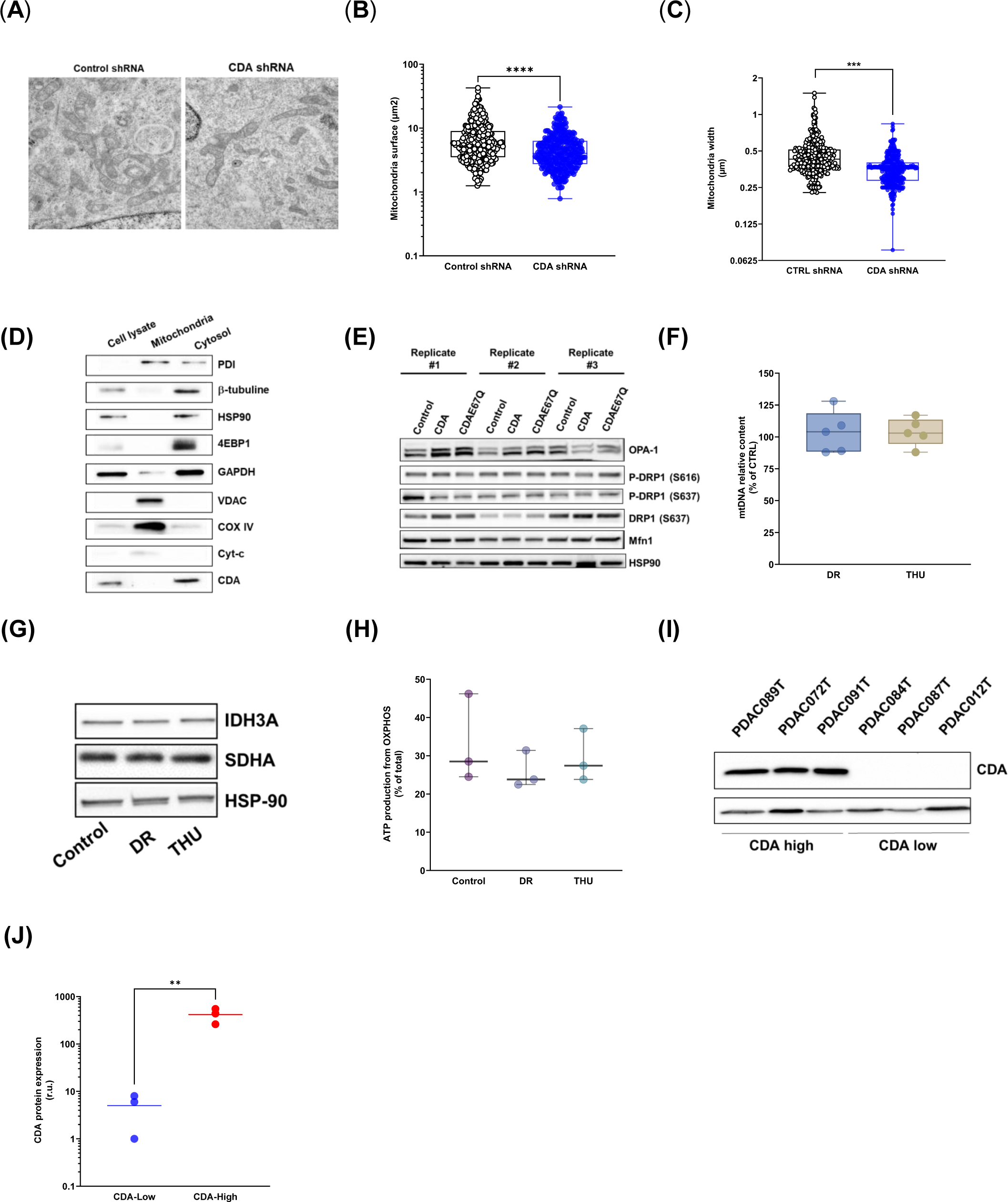
**A**. Representative captions of mitochondrial content using electron microscopy. **B**. Analysis mitochondrial surface in Mia PaCa-2 cells expressing shRNA against CDA. Results are presented as mean ± min to max of 330 mitochondria from two independent experiments using two different cell preparation, ****: p<0.001). **C**. Analysis of mitochondrial width. Results are presented as mean ± min to max of 330 mitochondria from two independent experiments using two different cell preparation, ***: p<0.005) in Mia PaCa-2 cells expressing shRNA against CDA. Cells expressing random shRNA were used as control. **D**. Western blot analysis of protein disulfide isomerase (PDI), β-tubulin, HSP90, 4EBP1, GAPDH, VDAC, COX IV, cytochrome C and CDA in cell lysate, mitochondrial fraction or cytosol of BxPC-3 PDAC cells that express high endogenous level of CDA. Representative of four experiments performed using two different cell preparation. **E**. Western blotting for OPA, DRP-1, phospho-DRP1 (P-DRP1), Mfn1 and CDA in Mia PaCa-2 cells overexpressing wtCDA or CDAE67Q. Cells expressing luciferase were used as control. Representative of three experiments performed using three different cell preparation. Western blotting for IDH3A and SDHA (**F**), mitochondrial DNA quantification by qPCR (**G**) and ATP production from OXPHOS (**H**) in Mia PaCa-2 cells treated by catalytic inhibitors of CDA (DR and THU). Results are representative of three independent experiments and expressed as mean ± min to max of five independent experiments analyzed in triplicate and are mean ± min to max of three independent experiments analyzed in triplicate. Western blot analysis of CDA expression (**I**) and corresponding quantification (**J**) in PDAC primary cells. Results are representative and expressed as mean of four independent experiments. **: *p*<0.01.

## ACKNOWLEDGMENTS

The authors thank the financial support of the “Fondation Toulouse Cancer Santé” (Audrey Frances), “Inserm” (Audrey Lumeau), “Région Occitanie” (Audrey Lumeau), “Université Paul Sabatier UT3” (Marion Gayral), “Fondation de France (n°00097692)” (Nicolas Béry) and “Ligue Nationale Contre le Cancer” (Audrey Lumeau). The authors thank Ms Emilie Martin and Catherine Zanibellato for technical assistance, and Drs. Sophie Vasseur (Cancer Research Centre of Marseille, France), Christian Frezza (Univ. Koeln, Germany) and Matthew Vander Heiden (Koch Institue, MIT) for helpful discussions. We are grateful to the MetaTOUL platform (Genotoul) for performing metabolomics experiments, and to the Imaging platform of University Paul Sabatier for performing electron microscopy.

## AUTHOR CONTRIBUTIONS

Conceptualization: Pierre Cordelier, Jean-Emmanuel Sarry, Anthony Lemarie. Investigations: Audrey Lumeau, Audrey Frances, Marion Gayral, Estelle Saland, Lucille Stuani, Delphine Pagan, Naima Hanoun. Supervision, Pierre Cordelier, Jean-Emmanuel Sarry, Jean-Charles Portais, Nicolas Béry, Louis Buscail, Jérôme Torrisani, Nelson Dusetti, Anthony Lemarie. Writing – original draft: Pierre Cordelier. writing – review & editing: Jean-Emmanuel Sarry, Nelson Dusetti, Nicolas Béry, Anthony Lemarie, Delphine Pagan, Audrey Frances. Funding acquisition: Pierre Cordelier, Jean-Emmanuel Sarry; Anthony Lemarie. Data analysis: Jean-Emmanuel Sarry, Audrey Lumeau, Pierre Cordelier, Nelson Dusetti, Antony Lemarie, Audrey Frances, Marion Gayral, Nicolas Béry, Jean-Charles Portais.

## MATERIALS AND METHODS

### Cellular models

Mia PaCa-2 cells expressing luciferase, wtCDA, CDAE67Q, control shRNA or shRNA targeting CDA have been described elsewhere^4^. Pancreatic cancer patient-derived primary cultures were cultured as described before^11^.

### Proliferation assays

Cell confluence was monitored non-invasively using the IncuCyte Zoom (Sartorius, Essen BioScience). Six thousand cells were seeded in 96-well plates in 100μl of complete medium for 24 h. Medium was changed and cells were treated as indicated. Cell confluence was analyzed using Zoom software version 2018B and expressed as area under curve (AUC).

### RNA extraction, RNAseq and functional enrichment

Total RNA was purified from cells using standard approach (miRNAeasy) following the manufacturer’s recommendations (Qiagen). Messenger RNA sequencing was performed by Novogene (Cambridge, UK). We performed enrichment analysis using GSEA Hallmarks and Reactome^15^ and most other pathways were extracted from MSigDB. No genes were removed during the analysis.

### Metabolic analysis

Five hundred thousand cells were seeded in 6-well plates with 2mL medium for 24h. Metabolites were quenched and extracted in a one-step process as described previously^16^. C^13^ IDMS standards added into the quenching solution were used to quantify metabolites, and samples were analyzed by the MetaTOUL platform (Toulouse).

### Mitochondrial DNA (mtDNA) quantification

Total cellular DNA was extracted with QIAamp DNA mini kit (Qiagen) according to the manufacturer’s instructions. DNA quality and quantity were measured using Nanodrop (Thermo Scientific). mtDNA and genomic DNA content were quantified by qPCR as previously described ^17^.

### Protein extraction and Western blot analysis

Cell pellets were incubated in Radio-Immunoprecipitation Assay buffer (RIPA, 150mM NaCl, 0.5% sodium deoxycholate, 0.1% SDS, 50 mM Tris-HCl, pH 8.0) supplemented with 10μL/mL of protease inhibitor (Sigma-Aldrich). After 15 min on ice, samples were centrifuged (15 minutes at 12 000x*g* and 4°C) and supernatants containing soluble proteins were collected. For cytoplasmic extract isolation, cell pellets were resuspended in 10mM Tris-HCl, pH 7.4 containing 1.5mM MgCl2, 5mM KCl, 0.5mM dithiothreitol, 0.5%NP40 and 0.5mM PMSF complemented with 10μL/mL protease inhibitors (Sigma-Aldrich) and incubated on ice for 10 min. Following centrifugation for 15 min at 2000x*g* and 4°C, the supernatant corresponding to the cytoplasmic fraction was collected. Mitochondrial protein isolation was performed as previously described^18^. The pellet containing the mitochondrial fraction was solubilized in RIPA buffer as previously described.

### Flow cytometry

Cells were collected by trypsinisation and resuspended in Phosphate-buffered saline and incubated with 200μM cyanide 3-chlorophenylhydrazone (CCCP, Sigma) for 15 min as TMRE negative control, 20μM MitoSox, 200nM tetramethylrhodamine, ethyl ester (TMRE, Sigma) or 100nM MitoTracker Green (MTG, Sigma) for 20min. Cells were resuspended in Annexin-V Binding Buffer (BD Pharmingen) and 1μL/mL anti-Annexin-V antibody (BD Pharmingen) was added to measure cell viability. Mitochondrial ROS levels were measured by flow cytometry using MitoSox (ThermoFisher), mitochondrial membrane potential and mitochondrial mass were measured with TMRE (ThermoFisher) and MTR (ThermoFisher) respectively, within Annexin-V negative, live cells fraction.

### Quantification of ATP production

ATP was measured using the Cell Titer Glo kit (Promega). Eight thousand cells were plated into a 96-well plate (Corning) for 24hours. Medium was changed and cells were incubated with 100μL of PBS as control, or treated with 100μM sodium iodoacetate (IA, Sigma) to block glycolytic ATP production, alone or in combination with 30μM Trifluoromethoxy carbonylcyanide phenylhydrazone (FCCP, Sigma) to block mitochondrial ATP production. Following 1-hour incubation, 100μL of Cell Titer Glo reaction mix solution were added to each well. Plates were then analyzed for luminescence with a Clariostar apparatus (BMG LABTECH). Percentage of glycolytic ATP = 100x(ATP_PBS_ – ATP_IA_) / (ATP_PBS/water_ – ATP_IA+FCCP_). Percentage of mitochondrial ATP = 100-% Glycolytic ATP.

### Quantification of oxygen consumption rate (OCR) and extracellular acidification rate (ECAR)

Cells were seeded at 4×10^4^ cells per well in Seahorse XF24 tissue culture plates (Seahorse Bioscience Europe). The sensor cartridge was placed into the calibration buffer medium supplied by Seahorse Biosciences to hydrate overnight. Twenty-four hours later, medium was changed with 500μL of XF base minimal DMEM medium containing 10mM glucose, 1mM pyruvate and 2mM glutamine at pH 7.4. After one-hour incubation at 37°C in CO2 free-atmosphere, basal oxygen consumption rate (OCR, as a mitochondrial respiration indicator) and extracellular acidification rate (ECAR, as a glycolysis indicator) were quantified using the XF analyzer, according to the manufacturer’s instructions. The data were normalized to protein content, determined by Bradford assay following cell lysis using 0.1N NaOH.

### Transmission electronic microscopy

Cells were fixed with 2% glutaraldehyde in 0.1 M Sorensen phosphate buffer (pH = 7.4), then washed with the Sorensen phosphate buffer (0.1 M) for 12 hours. Cells were incubated with 1% OsO4 in Sorensen phosphate buffer (Sorensen phosphate 0.05 M, glucose 0.25 M, OsO4 1%) for 1 hour, washed twice with distilled water and pre-stained with 2% uranyl acetate aqueous solution for 12 hours.

Samples were dehydrated in ascending ethanol solutions and embedded in epoxy resin (Epon 812, Electron Microscopy Sciences). After 24 hours of polymerisation at 60°C, ultrathin sections (70nm thick) were mounted on 150 mesh collodion-coated copper grids and post-stained with 3% uranyl acetate in 50% ethanol and with 8.5% lead citrate before being examined on a HT 7700 Hitachi electron microscope at an accelerating voltage of 80 KV. Number of mitochondria per cell and number of cristae per mitochondria were counted by blinded users without indication of experimental groups. Length and width of mitochondria was measured using the imageJ software.

### TCGA data analysis

HTSeq counts for PDAC were downloaded from the <u>GDC portal</u>. Curated PDAC samples were filtered based on information from ^13^. Samples were classified according to low (less than second quartile), normal (between second and third quartile) and high (higher than third quartile) values of CDA expression. Differential expression analysis comparing low CDA vs high CDA samples was performed with DESeq2 using low samples as the reference for the sign of the log2FoldChange.

### Statistical analysis

Unpaired Student’s *t*-or Wilcoxon-Mann-Whitney tests were used to determine the statistical significance of differences between two groups using GraphPad Prism 9 software with the default settings. Methods of statistical analysis are indicated in the figure captions. Values are presented as \**p*□<□0.05, \*\**p*□<□0.01 and \*\*\**p*□<□0.005 and **** *p*□<□0.001. Error bars are s.e.m. unless otherwise stated. No data were excluded from the analysis.

